# Ularcirc: Visualisation and enhanced analysis of circular RNAs via back and canonical forward splicing

**DOI:** 10.1101/318436

**Authors:** David T Humphreys, Nicolas Fossat, Patrick P L Tam, Joshua W K Ho

## Abstract

Circular RNAs (circRNAs) are a unique class of transcripts that can only be identified from sequence alignments spanning discordant junctions, commonly referred to as backsplice junctions (BSJ). The challenges of detecting a BSJ from short read high throughput sequencing (HTS) data has steered software development to focus primarily on algorithmic methods to accurately capture BSJs. Here we present Ularcirc, the first software tool that provides a complete circRNA workflow from detection, integrated visualization, quality filtering of BSJ and forward splicing junctions (FSJ), through to sequence retrieval and downstream functional analysis. More importantly, Ularcirc uses an innovative method to filter out false positive circRNAs coined read alignment distribution (RAD) score which allows detection of circRNAs independent of gene annotations. We used Ularcirc to characterise circRNAs from public and in-house generated data sets and demonstrate how to discover (i) novel splicing patterns of parental transcripts, (ii) internal splicing patterns of circRNA, and (iii) the complexity of BSJ formation. Furthermore, we identify circRNAs that have potential open reading frames longer than their linear sequence. Finally, we have identified and validated the presence of a novel class of circRNA generated from *ApoA4* transcripts whose BSJ derive from multiple sites within coding exons. Ularcirc can be accessed via https://github.com/VCCRI/Ularcirc.

## INTRODUCTION

High throughput sequencing is a technology regularly used to identify and quantitate transcripts that are present in enriched (polyA) or depleted (ribominus) RNA purifications. Circular RNAs (circRNAs) are a class of abundant transcripts that can be identified in most library preparations that are not geared towards poly(A) enrichment. The digestion of RNA with RNase R, which selectively degrades linear RNA [1], is currently the only method that is utilized to selectively enrich for intact circRNA transcripts [2]. However, with sufficient sequencing depth, circRNAs can be detected in standard ribosome depleted libraries [3-5]. Additionally, circRNAs can also be detected in specialized library preparations designed to capture specific RNA fractions (e.g., polysome, pulldowns).

CircRNAs are identified from sequence data that capture the junction of discordant exons, commonly referred to as a backsplice junction (BSJ). The BSJ comprises a canonical splice donor and an upstream splice acceptor which may be derived from the same exon (single exonic circRNA) or any upstream exon of a linear transcript (multi-exonic circRNA). Thousands of circRNAs have been identified from a wide range of animals and plants and catalogued in online databases [6, 7]. Many have been identified to be tissue specific and/or developmentally expressed. The lack of a 5’ or 3’ RNA termini provides significant stability to the circRNAs and their expression in diseases has found use as biomarkers [8].

Proposed mechanisms of circRNA biogenesis involve proximal localization of backsplice junctions via lariat formation or through base pairing of inverted complementary repeat sequences *[2*, 4, 9]. The later mechanism has been demonstrated through the use of mini transgenes [10] and is regulated post transcriptionally by ADAR enzymes which alter the nucleotide composition and thereby influencing the stability of hybridization [11]. A number of splicing factors are also capable of influencing circRNA formation, which include Quaking [12], Rbm20 [13] and Mbln [14, 15]. Quaking, an important factor in EMT, binds to motifs within intronic sequences and dimerises bringing backsplice junctions into close proximity [12]. The exact mechanism of how Mbln and Rbm20 regulate circRNA formation is yet to be elucidated but given that they influence promiscuous splicing patterns suggests they have the capability to influence intronic interactions either through lariat formation or possibly a motif based interaction.

The regulation of circRNA biogenesis and tissue specific expression patterns suggests that circRNAs are not by products of splicing events. Functional roles for circRNA are largely unknown but there is increasing evidence that some circRNAs can act as regulatory elements by decoying RNA or protein molecules. Specifically, circRNAs can bind miRNA through multiple complementary binding sites to one or more miRNAs [3, 16]. Similarly, sequencing motifs within a circRNA can also interact with RNA binding proteins sequestering them from other activity [14, 17].

CircRNAs have also been identified to be capable of being translated. In vitro models demonstrated that circRNAs with internal IRES elements are capable of translating reporter genes [12]. More recently ribosomal footprinting and mass spectrometry studies also support translation of circRNAs [18]. The absence of a cap and the poly(A) tail means circRNA translation must be non-canonical. N6-methyladenosine (m6A) residues are enriched in circular RNAs and m6A modifications has been shown to be a possible driver of cap-independent translation [19].

The growing interest in circRNA coincides with the development of specific bioinformatic algorithms that focus on detecting, quantifying and annotating reads that capture backsplice junctions. Recent reviews of circRNA software reveals that all algorithms are effectively capable of detecting BSJ but vary in their ability to filter out false positives [20]. However, to confidently identify and analyse circRNAs, further downstream analyses such as internal exon usage, comparison of BSJ to parental FSJ expression, ORF and binding motifs identification, as well as tools for splice junction visualization are necessary. At this juncture, these complementary analyses were commonly performed in-house using custom bioinformatic scripts. To our knowledge, there is only one publicly available tool that goes beyond the detection of BSJ to characterize the internal exon use of circRNAs [21]. There is therefore a void of software tools available to the community for comprehensive identification and analysis of circRNAs. With this in mind, we developed the software package: Ularcirc.

Ularcirc combines BSJ detection with other analytical tools: It quantifies BSJ in relation with forward splice junction (FSJ) and the usage of internal exons, and has an intuitive graphical user interface menu system allowing the user to navigate to genes/junctions of interest and ultimately generated dynamic integrated genomic visualizations of both BSJ and FSJ. Furthermore, Ularcirc provides preliminary functional analysis by identifying open reading frames and miRNA binding motifs. Ularcirc is an open source software package written in R language and built using a shiny framework (http://shiny.rstudio.com). Ularcirc is dependent on Bioconductor databases and packages enabling it to be easily setup for a range of conventional and emerging model organisms. Using Ularcirc, we analysed our own data and a number of public data sets to demonstrate the ability to discover novel processing and functional features of circRNAs.

## RESULTS

### Classification of reads aligned to circRNA

We classified four different sequence alignments for circRNA from data sets generated with current short read technologies and refer to these as Type I – IV **(Figure 1 A and B).** Type I alignments are those that cover nucleotides that span regions within a single exon or across multiple concordant exons. These nucleotides while within a circRNA are indistinguishable to positions of a linear RNA and cannot be used for identification. Type II and III are alignments that capture a BSJ in either the primary or paired end read respectively, and can be used as direct evidence of a circRNA existence. Type IV are discordant alignments of paired end reads that flank but do not contain a BSJ. On their own type IV alignments do not identify a BSJ and therefore should only be used as supporting evidence.

**Figure 1:**
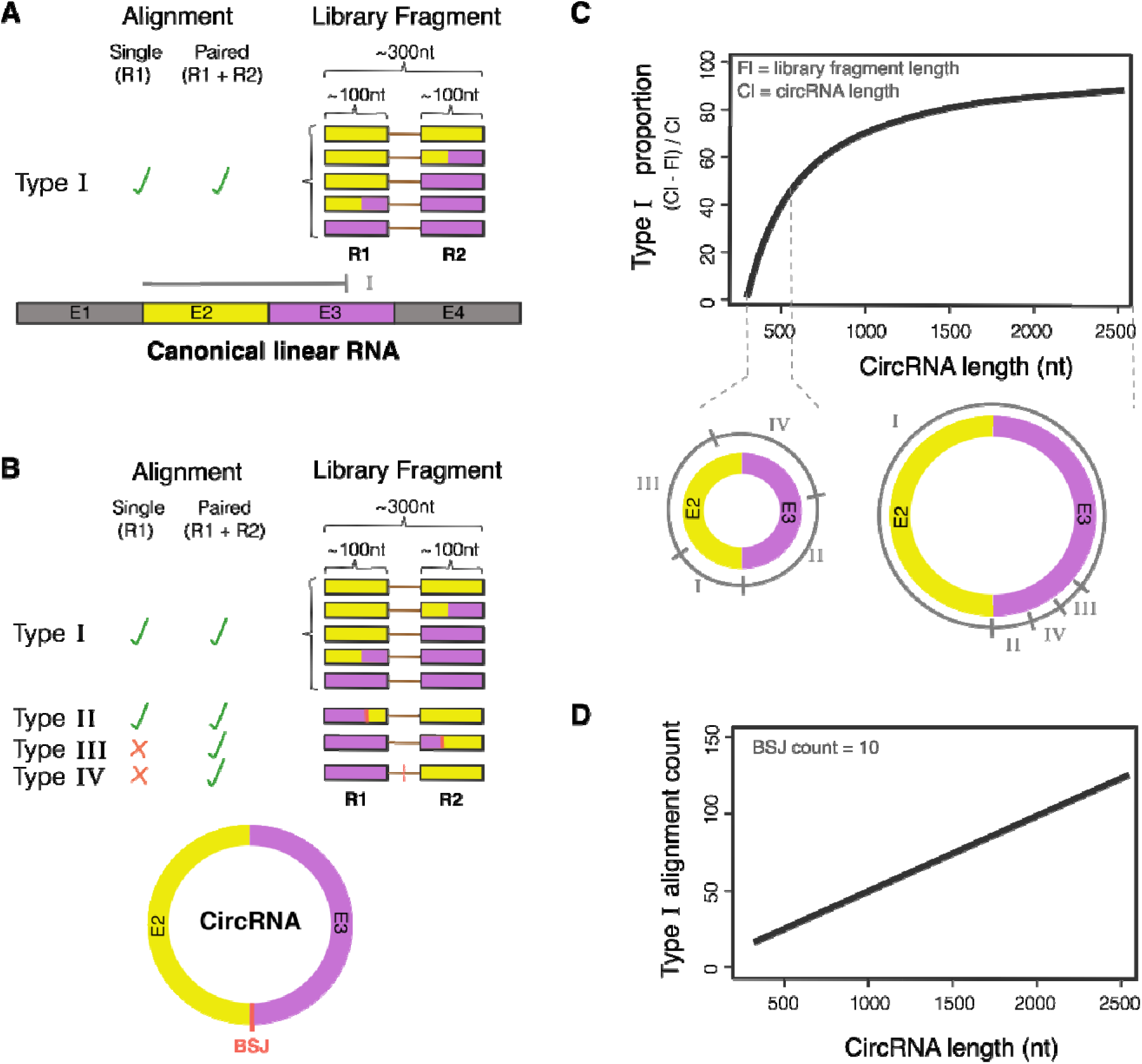
Theoretical modelling of sequence alignments from circRNA and linear RNA. A) Type I alignments are those that are indistinguishable between linear and circRNA. B) BSJ of circRNA are detected from type II and III alignments and can be inferred from type IV alignments. C) The proportion of type I alignments increase with circRNA size. The small and large circRNA schematics below the graph highlight the different proportion of alignment types across a small and large circRNA. The alignment ranges labelled I through to IV defines the 5’ start position of the aligned read pair. D) Inferred number of type I alignments from a different size range of circRNAs that have a theoretical BSJ count of 10.

We calculated the theoretical distribution of Type I – IV alignments and find that there is an increasing proportion of Type I alignments with circRNA size **(Figure 1C, materials and methods).** These calculations have been constructed on typical features of short read RNA-Seq libraries, where the average library fragment size is ^~^300bp and paired end read length is 100nt **(Figure 1B).** Consequently, the proportion of type I alignments is quasi null for circRNA smaller than a library fragment length (i.e 300nt). The proportion of type I alignments increases steeply in circRNA having sizes up to twice that of a generated library fragment (i.e. 600nt where type I alignments <=50%). This steep proportional increase is because type I alignments are becoming feasible in this size range **(Figure 1C).** CircRNAs that have a size greater than twice that of a library fragment (i.e. 600nt) result in a more gradual increase of type I-alignment proportion.

Type I alignments from a circRNA are reported in the standard BAM alignment files and cannot be distinguished from fragments generated from linear RNA. Therefore, in situations where a circRNA is abundantly expressed relative to linear parental transcripts (>10%), it is very likely that type I reads from the circRNA may contribute a significant proportion of aligned reads to the parental (pre-circularized/native) transcript, which may impact on various downstream analysis. To demonstrate this phenomenon, we calculated the theoretical number of type I alignments that are produced from different sized circRNAs with an arbitrary fixed absolute BSJ read count of 10. We found that the number of type I alignments will be multiples of the BSJ count as the size of a circRNA increases **(Figure 1D).** Ularcirc identifies and quantitates type II and type III junctions from sequencing data sets. Additionally, Ularcirc identifies type I FSJ alignments. Importantly, Ularcirc does not filter splice junctions on existing gene models and therefore can potentially discover novel junctions.

### The Ularcirc pipeline

BSJ and FSJ can be captured in RNA-sequencing data through reads covering discordant and canonical junctions respectively. Several aligners are capable of correctly aligning reads than span FSJ, but very few implement the detection of BSJ. The STAR aligner has become one of the more popular RNA-Seq aligners due to its speed and accuracy [22, 23]. However, it is less well known that the STAR aligner can also provide output count tables for both canonical and non-canonical “chimeric” junction counts. These count tables are simple tab delimited text files that are considerably smaller than traditional aligned BAM files and can be easily processed on desktop computers. We saw an opportunity to utilise this resource and designed Ularcirc to integrate FSJ and BSJ information by generating insightful visualizations, using the “Sushi” genomic visualization library [24], and build detailed annotated count tables for in depth analysis.

Ularcirc provides real time data analyses by dynamically generating an integrative genomic view (IGV) of BSJ and FSJ counts annotated with known transcript exon/intron boundaries. Junction count data is plotted as loops, where the loop height represents sequence counts and x-axis intersections represents the genomic coordinates of the splice junction. Junction data is overlaid onto a pre-existing gene model where transcript exons are graphed as rectangle blocks, similar to what is found in most genomic viewers **(Supplementary Figure 1A).** Ularcirc also provides a zoom functionality enabling resolution for dense and complex gene structures **(Supplementary Figure 1B).** Furthermore, any FSJ or BSJ can be selected from raw data tables and the corresponding splice junction in the IGV is colored.

Ularcirc is written in R using the shiny framework and the application can be launched as a standalone program on individual machines or as a multi-user setup through a server or cloud installation. Ularcirc has an intuitive graphical user interface while the backend automatically detects installed Bioconductor resource databases and other libraries **(Figure 2).** As of Bioconductor release 3.5 there are 13 organisms with complete genome, annotation and transcript databases **(Supplementary Table 1).** Ularcirc is immediately applicable for all these organisms once the appropriate libraries are installed. Furthermore, Bioconductor has workflows for the incorporation of other organism genome and annotation files, thereby making Ularcirc compatible for any organisms.

**Figure 2:**
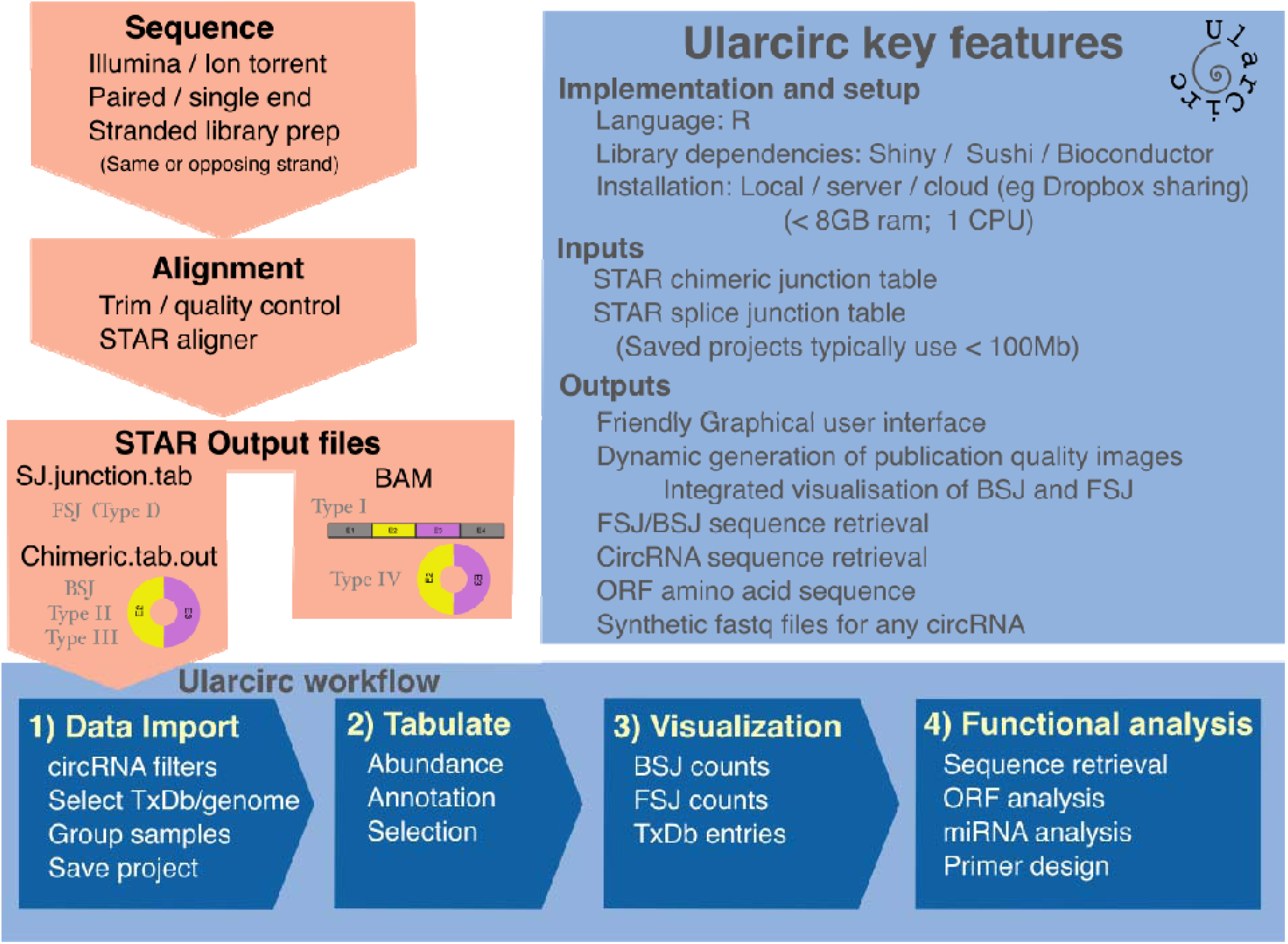
Schematic of Ularcirc software key features and workflow. Ularcirc requires two junction files from the STAR aligner and from this provides integrated visualisation and analysis of backsplice junctions (BSJ) and forward splice junctions.

### Annotating, quantifying and quality filtering of circRNA

We designed Ularcirc to follow a simple four-step workflow that enables quick identification of putative circRNAs **(Figure 2).** Using default parameters, users can generate a list of annotated abundant circRNAs from 1-10 data sets within minutes using a standard desktop computer. At this juncture, two additional annotation options are available. One option provides coverage information of parental linear FSJ that are internal, external or span over circRNA boundaries. We believe parental FSJ, identified by type I alignment, have been under-utilised in circRNA analysis and anticipate that this will provide unique opportunities in downstream filtering and analysis. Below we describe examples of novel FSJ patterns that occur with an increasing BSJ count.

The other and most innovative option is the calculation of a read alignment distribution (RAD) score. The RAD score is only applicable for paired end sequencing data as it is the ratio of type II vs type III alignments (RAD = type II/(type II + type III)). In theory, genuine circRNAs should generate equal proportions of type II and III alignments and therefore have a RAD ratio approaching 0.5 **(Figure 3A).** Reciprocally, RAD score that is close to either 0 or 1 represent candidates which are captured from either type II or type III alignments respectively. In these situations, the annotated BSJ is likely to be derived from misalignments of linear sequences that capture homologous sequences (as identified in [25]) and not genuine circRNAs **(Figure 3B).**

**Figure 3:**
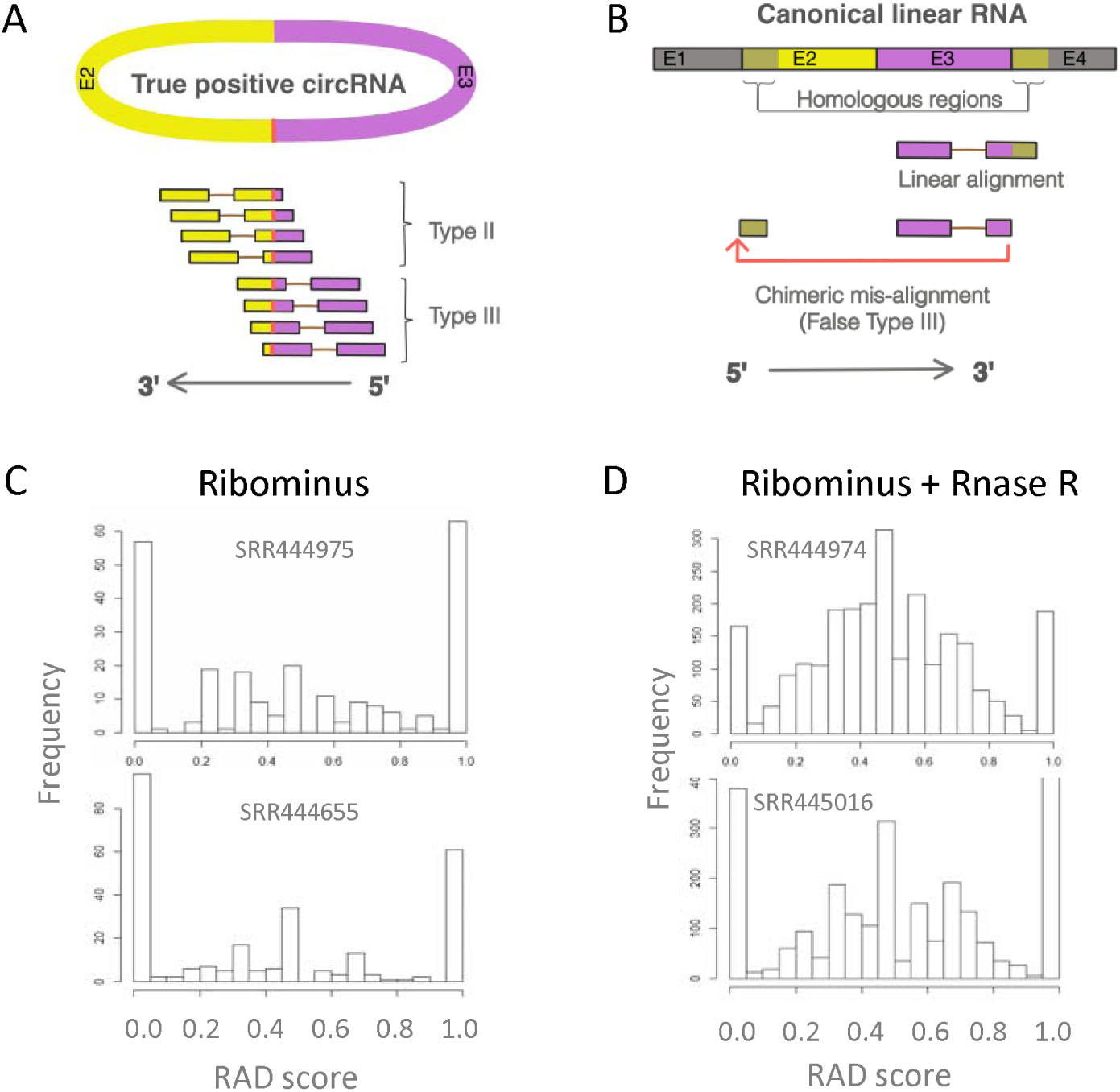
Alignment features that distinguish true and false positive circRNAs. A) True positive circRNAs display a spectrum of alignments that are captured on the primary or paired end read alignment and are referred to type II or type III respectively (refer Fig 1B). The RAD score for True positive circRNAs theoretically should be close to 0.5. B) Many false positive circRNAs are results of mis-alignment of linear reads to homologous regions within exons. The RAD score for these candidates approach a value of either 0 or 1. C) Ribominus data sets contain a mixture of true positive circRNAs and false positive circRNAs as defined by the RAD score. D) Rnase R data sets are significantly enriched for circRNAs with the majority of candidates having a RAD score approaching 0.5

To demonstrate the utility of this metric, we analysed the distribution of RAD scores from matched ribominus and ribominus+RNase R treated data sets **(Figure 3C and D respectively).** In all datasets, we noticed an abundance of BSJ having a RAD score very close to either 0 or 1, followed by a broader spectrum of RAD scores between 0.2 to 0.8. It is likely that BSJ producing RAD scores of 0 or 1 are false positives as these values become less abundant relative to the values of BSJ with RAD scores between 0.2 to 0.8 when the datasets are treated with RNase R. In support of this point, we noticed that a prominent false positive BSJ derived from adjacent myosin genes, previously examined from heart data sets [25], produced a RAD score of 1. True positive BSJ, which are more abundant in RNase R data sets, displayed an enrichment of RAD scores approaching 0.5 **(Figure 3D).** The broad deviation of RAD scores surrounding 0.5 may reflect the uneven distribution of RNA-Seq library fragments and/or detection biases in detecting the BSJ. Given this possibility, we recommend applying a strict threshold to identify and filter out false positives which conversely implies a loose threshold on retaining putative BSJ. Currently, the outcome filters out any BSJ (from paired end data sets) that has a RAD score <0.05 or > 0.95. Later in this manuscript we compared the output of this RAD filter setting to other software implementations.

### Visualisation and analysis of circular and linear transcripts

Ularcirc has functionality to display and analyse transcript sequences from both circRNA and linear transcripts through the integration of Bioconductor transcript coordinate and genomic databases. CircRNA sequences are generated by concatenating parental transcript exons sequences within the boundary of a selected BSJ. Users can select to display junction sequences from either a BSJ or FSJ, which enables easy primer design or downstream analysis. Additionally, Ularcirc has two functional analysis tools that can be used on a selected circRNA, which are (1) identification of the longest open reading frame (ORF) and (2) miRNA binding motif analysis.

The open reading frame analysis tool provides a circos-like plot representation of the longest open reading frame detected within a selected circRNA. The amino acid sequence derived from the ORF is also displayed enabling downstream analysis. It is known that some circRNAs contain long ORF, for example the circRNA from Slc8a1 contains a long ORF that has been tentatively linked to a truncated translated product [26]. Using Ularcirc, we identified a number of other abundant cardiac circRNAs encoding significantly long ORF. Interestingly we identified circRNAs where the ORF extends across the full circle past the initial start codon **(Supplementary Figure 2).** For example, the ORF identified within a circRNA derived from HipK3, a single circularized exon (1099 nt in size), can in theory produce a 388 amino acids product which is 22 amino acids longer than what is possible on an equivalent length linear transcript. Interestingly the BSJ from Hipk3 circRNA is one of the most abundant in the developing heart and has a similar count to canonical FSJ of the linear parental construct.

The other function analysis tool within Ularcirc identifies miRNA binding sites. Ularcirc provides functionality to visualise the locations of miRNA binding sites of a selected circRNA (refer methods). Target sites are identified from miRNA seed sequences whose start position and size can be defined by user. The limited complexity in a short miRNA seed sequence can result in many single hits. Ularcirc provides the capability of filtering miRNA binding sites that occurs at a minimum frequency. After filtering, the short-listed binding sites are presented as a circos-like plot and the specific miRNA and the number of binding sites tabulated **(Supplementary Figure 3).**

### Splicing analysis of linear transcript and circRNA biogenesis

We wanted to assess if the FSJ output generated by the STAR aligner was capable of reproducing known alternative splicing patterns. In the heart, a number of alternative splicing events occur between postnatal days 1 and 28 as described by Giudice et al., 2014 [27], RNA-Seq datasets from this study were aligned with STAR and the junction output files were imported into Ularcirc. Ularcirc was clearly capable of visualising the alternative splicing patterns between postnatal day 1 and adult tissue **(Supplementary Figure 4)** and we therefore concluded that the FSJ output files are a valuable resource in determining exon usage patterns of parental transcripts.

The visualization and quantification of both FSJ and BSJ provide a unique opportunity to cross examine parental transcript expression with circRNA biogenesis. FSJs that span/splice across a BSJ potentially discriminate two previous models proposed for circRNA biogenesis **(Supplementary Figure 5).** The presence of a FSJ that spans a BSJ would be supporting evidence for lariat derived circRNA biogenesis (model 1 of [4]) resulting from exon skipping. Conversely the absence of spanning FSJ suggests circRNA biogenesis that is independent of exon skipping. From all data sets analysed **(Supplementary Table 2)** there was a distinct absence of FSJ that span BSJ, even when the proportion of BSJ relative to other FSJ was greater than 10% **(Supplementary Figure 6).** One interpretation of this result is that the majority of circRNAs are actively produced and not by-products of lariats generated during canonical splicing. This was a similar conclusion to the study performed by Aufiero et al [28]. However as standard RNA-seq does not reliably capture FSJ from unstable linear transcripts we cannot rule out that circRNA biogenesis may also result from aberrant transcription and splicing events as reported in [29].

We next used Ularcirc to analyse expression patterns of abundant circRNAs and FSJ from parental transcripts across embryonic heart development data sets [25]. Interestingly we identified two genes that express abundant circRNAs, *Hipk3* and *Slc8a1*, whose expression correlated with novel parental canonical FSJ (red splice junctions in **Figure 4A and B).** For both *Hipk3* and *Slc8a1*, the novel FSJ was either upstream or downstream of the circRNA respectively. The correlation in circRNA and novel FSJ abundance suggests that for these examples, novel transcript isoforms produce the circRNAs.

**Figure 4:**
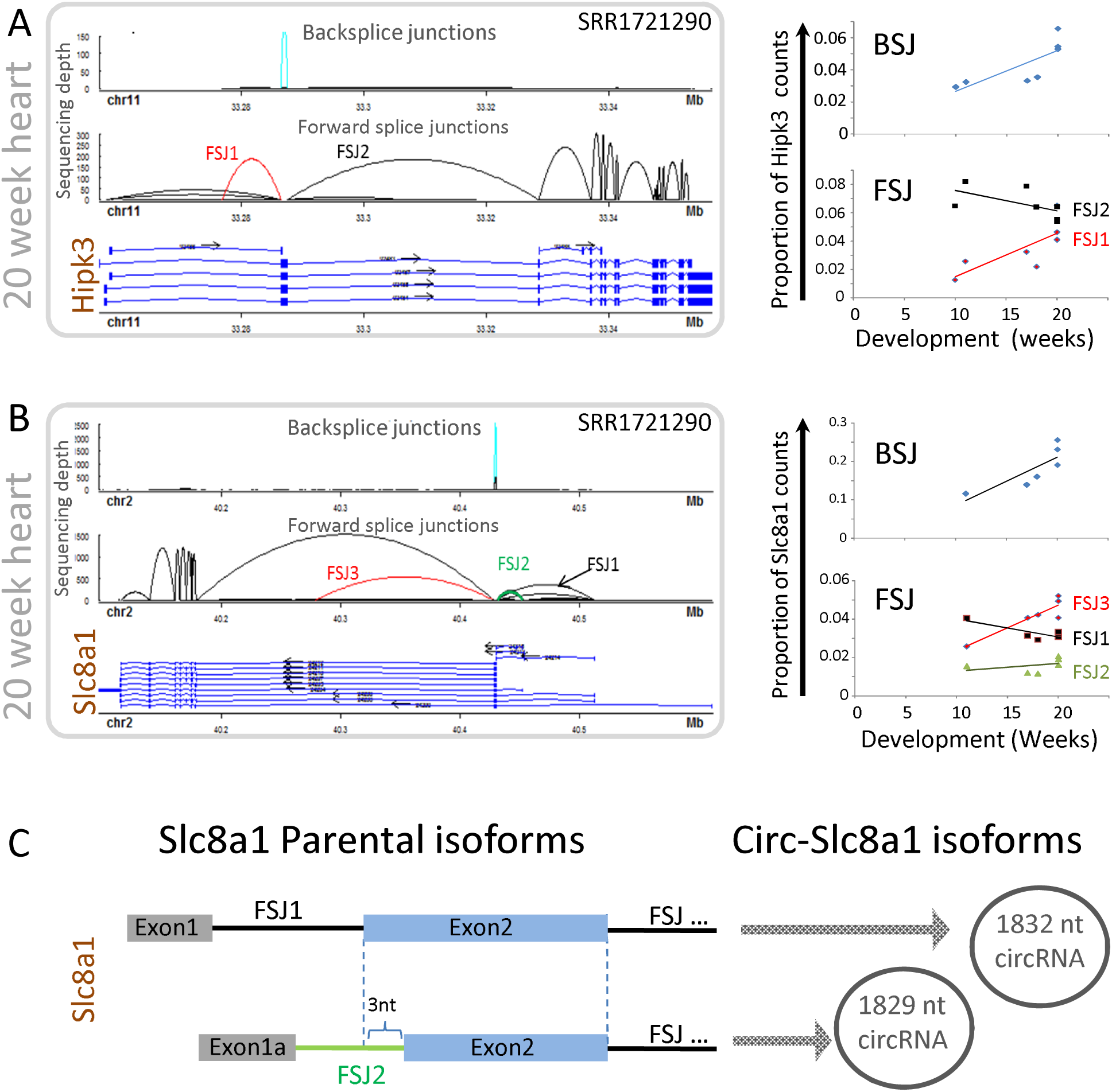
Examples of unique forward splice junction (FSJ) expression across development that is associated with one set of BSJ (circRNA) formation. A) circRNA from HipK3 is derived from exon 2 (light blue) whose expression correlates with a novel canonical FSJ of the parental gene (red). B) Two circRNAs from Slc8a1 are derived from exon 2 (previously reported as exon 1) that correlate with different exon expression patterns of the parental transcript. C) Each Slc8a1 circRNA is likely to derive from parental transcript isoforms that utilise different exon 2 splicing acceptor that differ by 3 nt.

Human *Slc8a1* produces two abundant circRNA isoforms that are upregulated throughout heart development. Previous analysis of *Slc8a1* circRNAs concluded that both circRNA isoforms are derived from alternative processing of exon 1 [25], which we thought may be unlikely. We therefore examined different gene models for *Slc8a1*. Refseq **(2017-07-13 annotation release)** annotates five Slc8a1 transcript isoforms of which only one contains an exon positioned upstream of the circularized exon. Genecode (v24) annotates eight Slc8a1 transcript isoforms of which four have an exon positioned upstream of the circularized exon **(Figure 4B).** Ularcirc visualization identified that heart Slc8a1 linear transcripts isoforms incorporate these alternative upstream exons. Therefore *Slc8a1*-circRNA should be annotated as deriving from exon 2. Furthermore, two of the dominant alternative upstream exons utilise distinct acceptor sites within exon 2 (FSJ1 and FSJ2 in **Figure 4B).** These different acceptor sites match perfectly to the BSJ of the two Slc8a1 circRNA isoforms. FSJ2 aligns to minor circRNA and FSJ1 aligns to major circRNA isoform **(Figure 4C).** Interestingly the novel linear and circRNA Slc8a1 transcript isoforms are not evolutionary conserved as the analysis of a mouse ventricle time course data set only identified one circRNA species that correlates with a single transcript isoform **(Supplementary figure 7).** However, mouse Slc8a1 circRNA are also derived from exon 2 supporting the notion that Slc8a1 circRNAs should be annotated as being processed from exon2 of the parental transcript isoforms.

### Internal circRNA splicing

CircRNAs are typically derived from one or more internal exons of parental transcripts. Deciphering internal exon composition typically involves specialized algorithms and/or long-read sequencing analysis [21]. We identified two scenarios where Ularcirc can also provide insights into the internal splicing patterns of circRNA, without the need for long-read sequencing. The first scenario involves looking for upregulated FSJ (FSJ hotspot), that resides within the coordinates of an abundant BSJ, relative to the other FSJ of the parental transcript. Ideally, the expression changes of a FSJ hotspot should correlate with the expression changes of the BSJ. Ribominus developmental time course data sets are a great resource for this type of analysis, particularly for developmentally upregulated circRNAs. One such example is circRNAs derived across five central exons of PTK2 in the developing heart. The FSJ within the BSJ boundaries increasing more than external FSJ suggesting they are derived from PTK2 circRNAs **(Figure 5A).**

**Figure 5:**
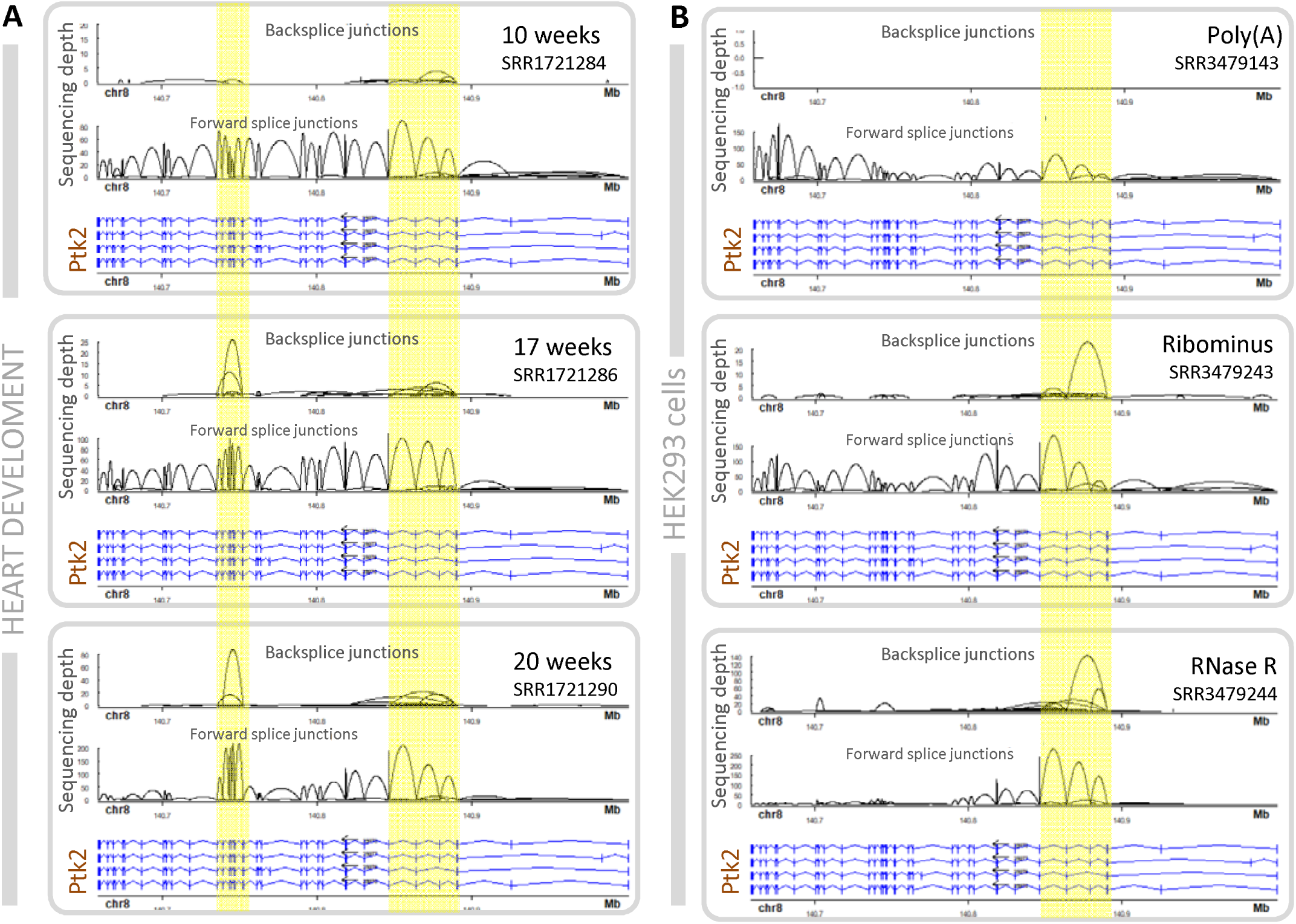
Ptk2 circRNAs in either human heart and Hek293 cells. A) Heart PTK2 circRNAs are mainly derived from the 5’ exons of parental transcript. Expression levels of FSJ (highlighted in yellow) within the boundaries of BSJs increase across development suggesting internal circRNA splicing. B) PTK2 circRNAs are mainly derived from the 3’ exons of parental transcript expressed in the Hs68 cell line and are detected in Ribominus and RNase-R libraries. FSJ within boundaries of BSJ (highlighted in yellow) are significantly enriched after RNase R treatment, suggesting internal circRNA splicing.

The second scenario also involves identifying FSJ hotspots that correlate with BSJ expression but utilizes data sets generated from RNase R treated RNA. We examined matched poly(A), ribominus, and RNase R public data sets generated from the HEK293 human cell line. As expected looking at *PTK2*, the circRNA isoforms detected by the BSJ in the ribominus data are enriched in the RNase R dataset but are absent, as visualized by the lack of BSJ, in the poly(A) data. Concomitantly, the FSJ within BSJ boundaries are lower in the poly(A) dataset compared to the ribominus dataset while counts of all FSJ junctions across the length of the rest of the gene are similar between the two datasets. Reciprocally, the only FSJ detected in the RNase R data are within the boundaries of the BSJ, which again supports that the exons within FSJ boundaries enter the composition of *PTK2* circRNAs **(Figure 5B).** We believe algorithms could be developed to take advantage of this trend and extrapolate internal splice junction composition of circRNAs. Furthermore, these two examples of circRNAs from the same gene **(Figure 5A and B)** also reveal that different tissues are capable of producing different circRNA isoforms. CircRNAs derived from the 3’ end of PTK2 are predominantly processed in Hs68 fibroblast cells, while circRNAs derived from the 5’ end are predominantly processed in heart tissue. This suggests that certain tissue-specific factors can drive circRNA biogenesis.

### Complex circRNA formation

Ularcirc can also capture the formation of more complex circRNA. Using a dataset generated in-house from murine epithelial cells of the small intestine, we identified an extreme example of complex circRNA formation derived from *ApoA4. ApoA4* encodes the secreted Apolipoprotein A4 which is important for lipid metabolism [30]. These circRNAs are not processed from defined linear acceptor/donor splice sites but rather a more disorganized spread of positions located within exonic regions **(Figure 6A).** We reasoned that if these BSJ represented genuine circRNAs, the majority of the BSJ could be validated in a PCR reaction using a single pair of divergent primers. Indeed, PCR amplification with this pair of primers produced a range of products that was represented by a smear on an agarose gel with a spread of larger incremental fragments **(Figure 6B and C).** The PCR product was then prepared as a sequencing library using fusion primers (see material and methods) and the resulting reads fully reproduced what we identified from the original RNA-Seq data **(Figure 6D).** To further confirm circularization, we replaced one of the divergent primer of the divergent pair with a primer that overlaps one defined BSJ and should therefore specifically amplify one type of circRNA. Visualization of the result of this PCR on agarose gel **(Figure 6E)** shows products where the larger bands are concatemers of smaller fragments produced from rolling circle amplification from the reverse transcriptase, further proving the event of RNA circularization [31] that was confirmed by sanger sequencing of the cloned fragments extracted from the gel **(Figure 6F).** Finally, we demonstrated that *ApoA4* RNA circularization is conserved in human by performing *ApoA4* divergent RT-PCR using RNA sample from human small intestine **(Figure 6G).**

**Figure 6:**
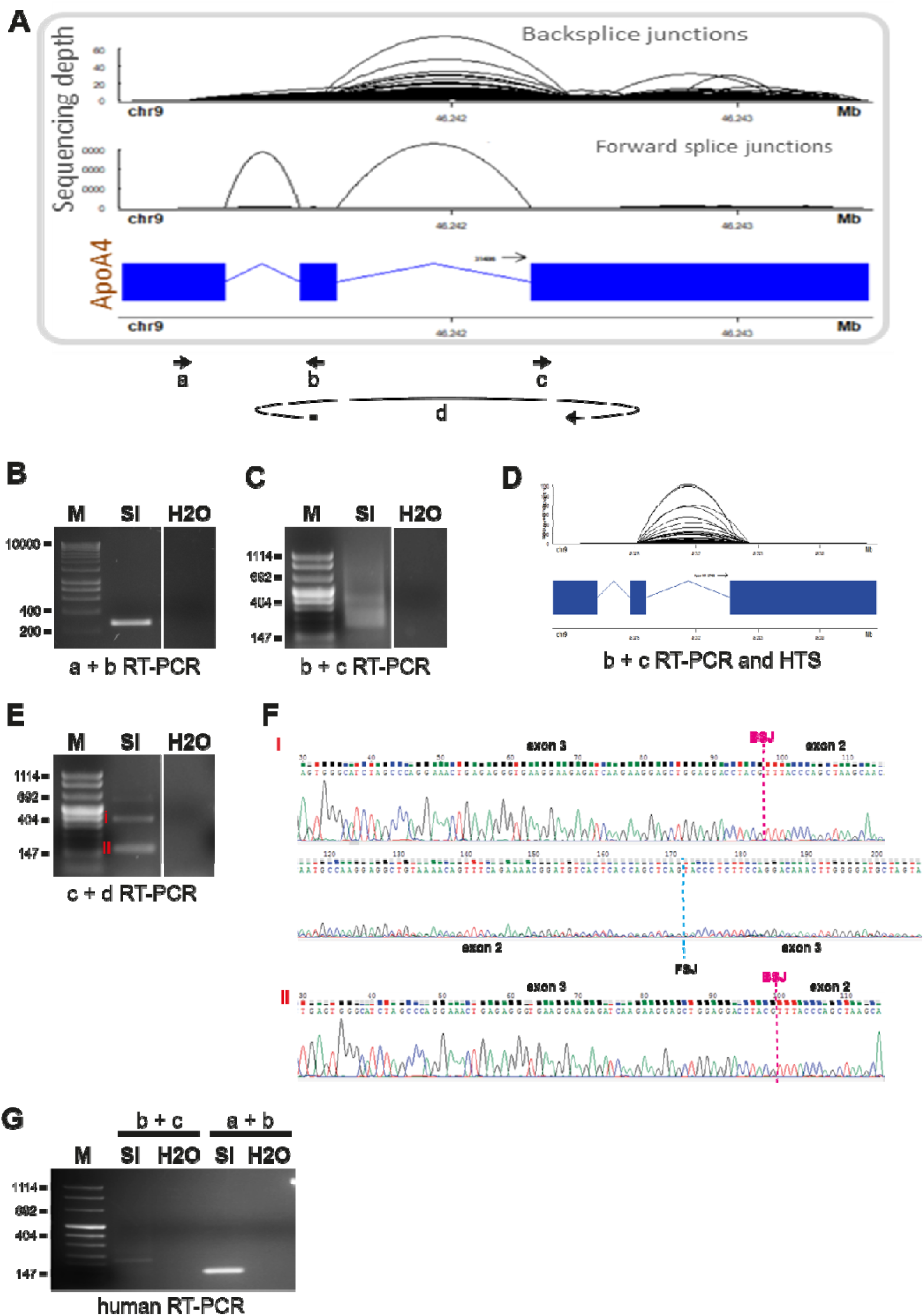
Complex circRNA formation from the *ApoA4* gene. A) High throughput sequencing and UlarCirc analysis of total RNA extracted from epithelial cells of the murine small intestine identified many BSJs derived from multiple positions within exons 2 and 3 of the *ApoA4* gene, as visualised with UlarCirc. Bold arrows indicate the position and the orientation of the primers used for PCR amplifications part B, C, D, E and G. Primer d overlaps a specific BSJ. B) Agarose gel visualisation of the parental (linear) ApoA4 transcript amplified by RT-PCR from total RNA with convergent primer a and b. C) Visualisation of multiple backsplice transcripts (smear) amplified with divergent primer b and c. D) High throughput sequencing of the products amplified with divergent primers b and c captures the same range of BSJs as seen in part A. E) Visualisation of a range of concatenated circRNA products (i.e. rolling circle amplification) amplified with divergent primers c and d (d overlapping a specific BSJ). F) Sanger sequencing of RT-PCR products (i) and (ii) isolated from the agarose gel part E. Positions of the BSJ and the FSJ are indicated. G) Visualisation of the backsplice and the parental (linear) *APOA4* transcripts amplified with divergent primers b and c and convergent primers a and b respectively, from human small intestine RNA. H2O, water-only control; HTS, high throughput sequencing; M, marker (size indicated on the left of the gels); PCR, polymerase-chain reaction; SI, small intestine.

While these observations further emphasize the potential complexity of circRNA formation, they show for the first time that BSJ can take place between RNA extremities that are not canonical acceptor and donor splice sites, which furthermore support the role of RNA binding proteins that are not known to be classical splicing factors for this process.

### Comparing Ularcirc to circExplorer2

The bioinformatics challenges in detecting circRNA from short-read sequencing datasets has engendered software development to improve the detection of BSJs and discriminate true from false positives. A review of five different software packages identified that each was capable of detecting a common pool of circRNAs, but each software was also capable of detecting a unique set of true positive BSJs [20]. The software package CircExplorer was considered as one of the best tools as it had high accuracy, good sensitivity and low memory consumption [20]. Furthermore, its successor, circExplorer2 can take inputs from a number of different aligners, including STAR, making it very versatile. We reanalysed data sets from Hs68 human fibroblasts [4] by aligning with the STAR aligner and compared the outputs of Ularcirc and circExplorer2 and found a high correlation in BSJ counts retrieved by both programs **(Figure 7A).** All abundant BSJ identified by circExplorer were also identified by Ularcirc with the majority having identical counts (correlation = 0.916). This comparison also demonstrates that the default inbuilt RAD thesholds filter in Ularcirc does not filter out true circRNAs. Interestingly, a number of novel BSJ were identified specifically by Ularcirc. Closer examination revealed that circExplorer2 failed to identify some of these candidates because they did not overlap with known gene transcripts splice junctions, a known limitation of circExplorer. To demonstrate circExplorers dependency and limitations on using known gene models we performed a theoretical exercise by re-analysing HEK293 RNase R treated RNA-Seq data sets [29]. We focused on ASAP1 gene which produces two alternative circRNA isoforms that use a common exon acceptor, as visualized with Ularcirc **(Figure 7B).** We modified the ASAP1 gene entry so that either the donor or acceptor exons that define a BSJ were shifted to either the 5’ or 3’ direction. Unsurprisingly circRNA were not reported by circExplorer2 when either donor/acceptor exon was modified **(Figure 7C).** Ularcirc however was able to identify both circRNAs without a gene reference.

**Figure 7:**
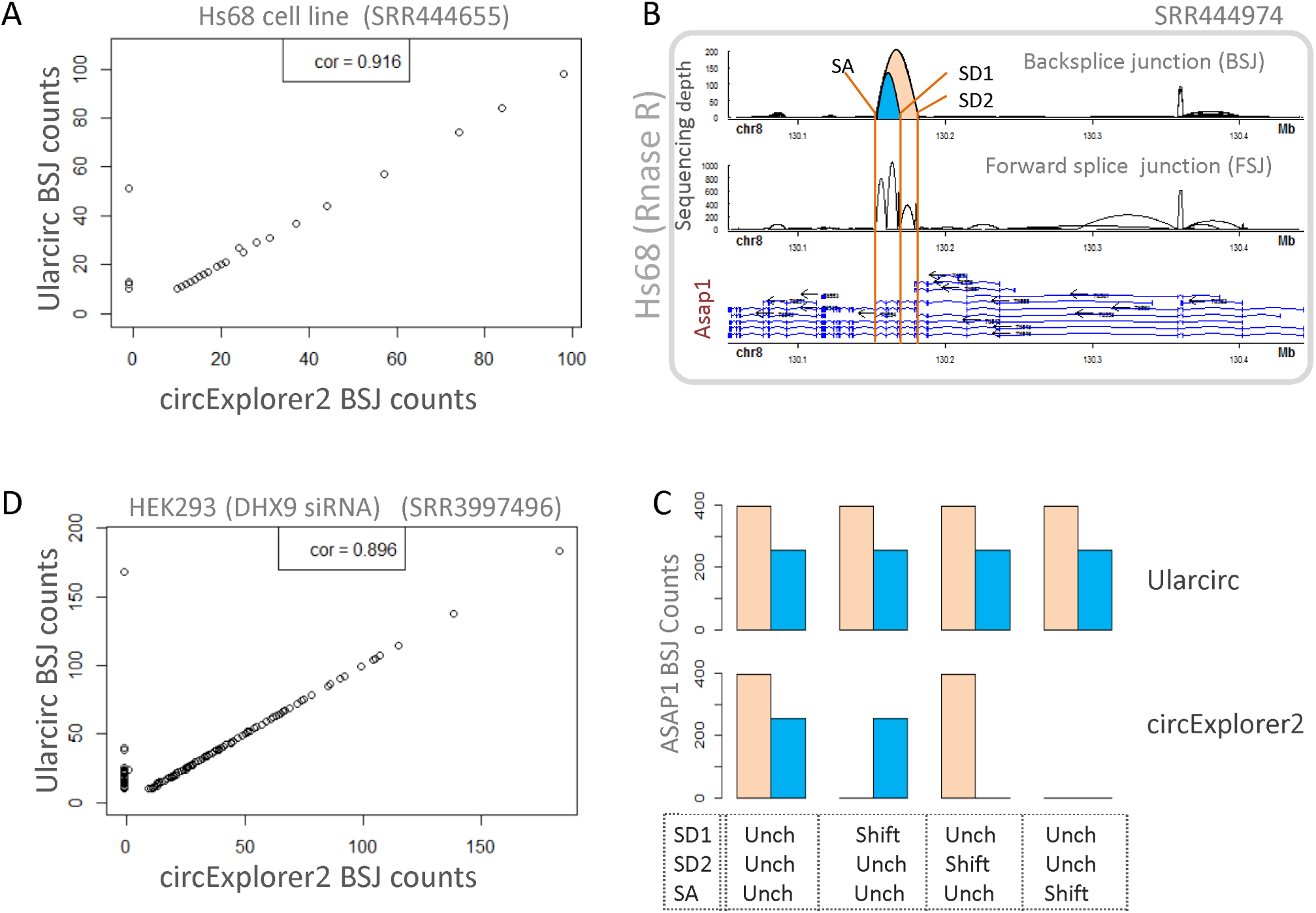
Ularcirc identification of BSJ is superior to circExplorer2. A) Comparison of BSJ raw counts recovered from Ularcirc and circExplorer2. B) Two Asap1 circRNAs share a common splice acceptor (SA) but originate from two different splice donors (SD1, SD2). C) Demonstrating the limitation of using gene model-dependent circRNA discovery. Gene entries for *ASAP1* were either left unchanges (Unch) or the splice sites listed in (B) were shifted 5’ and 3’ (shift) by 10nt before running the various software. CircExplorer2 fails to detect the circRNAs when corresponding *ASAP1* gene exons boundaries are shifted. Ularcirc identify both circRNAs even if the gene models are altered. D) Ularcirc identifies abundant circRNAs from *DHX9* knockdown cells which were previously missed due to the use of circExplorer2.

The use of circExplorer2 may result in some circRNA not being identified on genes that are not completely annotated. We re-analysed the circRNAs associated with knockdown of *DHX9*, which results in aberrant splicing of linear transcripts and the upregulation of many circRNAs. We discovered abundant circRNAs that were identified by Ularcirc (and having balanced RAD scores) but not by circExplorer2 **(Figure 7D).** As expected most of these circRNAs were derived from exon boundaries not defined in Gencode (release v24). For example, two circRNAs were identified to be processed from the 5’ end of Zranb1. Both ZranB1 circRNAs utilised the splice donor from exon 1 but create a BSJ either within the middle of exon1 or ~3kb upstream of exon 1 donor splice junction **(Figure 8A).** The analysis of FSJ expression patterns showed that ZranB1 has two alternative start sites that are not currently annotated, which we name 1B and 1C (pre-existing exon 1 now referred to as 1A). Furthermore, exon 1B and 1C splice into exon 1A to a splice acceptor position that exists within the current defined boundaries of 1A. The internal splice site within exon 1A matches the position of one circRNA **(Figure 8A inset).**

**Figure 8:**
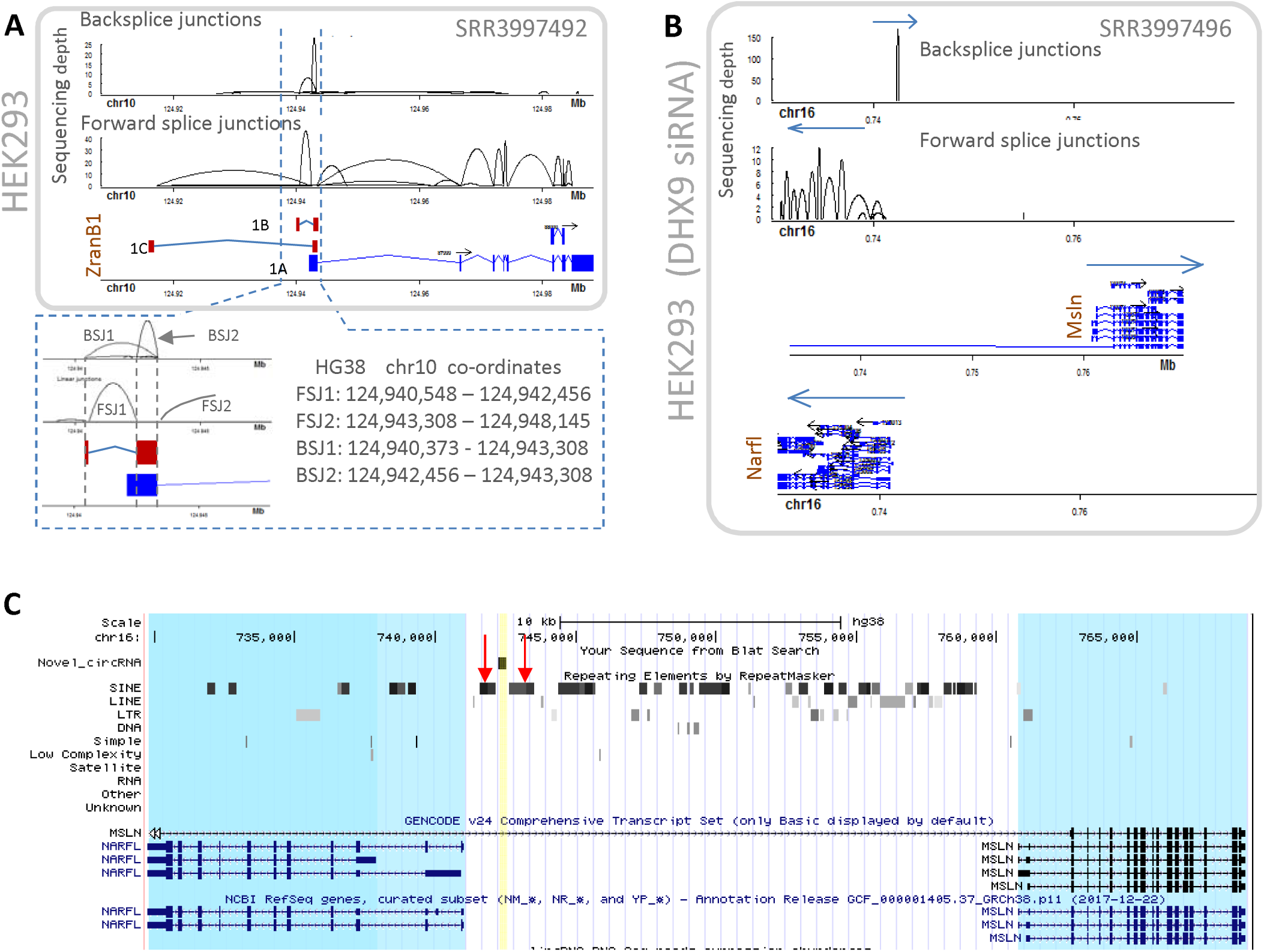
Novel abundant circRNAs that do not align to current gene annotations. A) We used Ularcirc to identify that ZranB1 has two additional alternative first exons which are annotated as 1B and 1C, which are both expressed in HEK293 cells. The pre-existing annotation is labelled as 1A Two circRNA isoforms are generated from transcript isoforms 1B and 1C as the coordinates of these backsplice junctions match perfectly to the novel transcript isoforms (refer zoomed inset). B) The most abundant circRNA from *DHX9* knockdown cells was identified using Ularcirc and does not align to any canonical annotated or expressed exon. Two neighboring genes that are transcribed from different strands are *NARFL* and *MSLN*. The novel circRNA is transcribed from the same strand as *MSLN*, but from an intronic region. C) UCSC browser screenshot highlighting the proximity of ALU repeat elements (red arrows) that surround the novel circRNA (highlighted in yellow). Neighbouring genes are highlighted in blue.

Ularcirc also identified an abundant BSJ that was annotated to be derived from a sub-intronic region of MSLN **(Figure 8B).** This circRNA is not flanked by annotated exon junctions but does have canonical splice donor and acceptor sites. However, there was no adjacent novel FSJ identified from the sequencing data. Interestingly, this circRNA was only expressed when *DHX9* is knockdown, suggesting that the circularization process is dependent on an RNA secondary structure. Analysis of the surrounding genomic landscape reveals an enrichment of SINE/ALU sequences that flank each side of the circularized sequence **(Figure 8C),** suggesting these sequences may contribute to circRNA biogenesis. The lack of associated FSJ and linear reads makes it difficult to identify the “parental” transcript. The two most likely possibilities are that the circRNA is derived from an antisense transcript from the NARFL promoter or from the spliced intronic sequence of the MSLN gene.

## DISCUSSION AND CONCLUSION

In this study we developed a bioinfomatic software package dedicated for circRNA detection, visualization and analysis. Ularcirc is built on the shiny - R framework and utilizes splicing junctions reported by the STAR aligner. We designed Ularcirc that follows a systematic workflow involving the generation of tabulated BSJ count tables, integrated genomic visualizations and functional analysis of ORFs and miRNA binding site. The graphical user interface offers investigators with minimal working knowledge in Bioinformatics the capability to explore the data sets. Furthermore, the integration of bioconductor databases allows the configuration of Ularcirc to render it compatible with the analysis of sequencing data of a wide range of organisms.

Existing circRNA software that focuses on the detection of BSJ is proven difficult to navigate, given the low frequency of BSJ in the data sets. One of the key challenges is the reliable filtering of false positive candidates that are predominantly derived from homologous exons [25]. By theoretical modelling, we predicted that an equal ratio of BSJ can be detected in both read pairs of paired end sequencing data. Based on this notion, we developed a read alignment distribution (RAD) score that can successfully identify false positive BSJs. This unbiased filter constitutes an innovative way to improve the accuracy of circRNA identification. The RAD score is a simple calculation that only requires minimal computational resources (Ularcirc calculates RAD score from CIGAR strings) therefore making its implementation very efficient. This method could prove useful in the future for applications such as the detection of fusion circRNAs derived from chromosomal translocations [32].

Currently, there is a “lack-of/weakness-in” the means to normalized BSJ counts. One approach is to normalize against total BSJ count, but given the wide discrepancies in circRNA discovery between software, the accuracy of this method will vary. Another approach is to normalize against the total linear reads (spliced reads per billion mapping = SRPBM) [4]) or a pre-defined set of canonical genes [12]. Ularcirc provides capability to build tables of counts of BSJ and associated FSJ (internal to the circRNA and external to the parental transcript). While evaluation of the best normalization method is out of the scope of the present study, Ularcirc does provide options to explore normalization of BSJ to parental transcript FSJ counts, which may be useful for analyzing abundant circRNAs.

We used Ularcirc to analyse BSJ and FSJ of circRNAs that had upregulated expression during development. This analysis identified examples of circRNAs whose increased expression coincided with an increased proportion of FSJs within circRNA boundaries. It is likely that this correlation may reflect internal circRNA exon usage. We further demonstrated that internal circRNA splicing could be inferred from visualising FSJ of RNAse-R digested RNA. Understanding the internal processing of a circRNA will provide the exact sequence content of that circRNA, and will be valuable for downstream functional analysis. A recent software package has been published to specifically interrogate circRNA internal splicing patterns and utilise algorithms that identify library fragment alignments that contain both a BSJ and a FSJ [21]. While this is an astute method to detect internal circRNA splicing, it is constrained to only detect FSJ within the library fragment length of a BSJ. We believe the visualization of internal splicing provided by Ularcirc highlights that new algorithms could be developed to compare RNase-R and time-course data sets to predict the complete internal splicing of long circRNAs. Our analysis of public data sets using Ularcirc has discovered novel FSJ within parental genes that flank circRNA exons. These novel exons potentially identify new parental transcript isoforms from which circRNAs are generated. This seems particularly likely for Slc8a1 as the novel FSJ that are positioned 5’ and 3’ of the circRNAs align perfectly with the coordinates of the minor circRNA isoform. Potentially, other similar cases will be revealed using Ularcirc.

CircRNA biogenesis competes with splicing machinery [13, 14, 33]. One hypothesis for circRNA biogenesis is that circRNAs are by-products of exon skipping [4]. If circRNA are indeed by products of exon skipping, we would expect to visualize the FSJ that span from the 5’ upstream exon through to the downstream 3’ exon. These splice junctions should be more apparent at the onset of circRNA biogenesis. However, we rarely detected alternative splicing events that could produce circRNA by products. Furthermore, we identified examples of circRNAs that are not derived from canonical splice junctions. In the case of *ApoA4* circRNAs, we could not detect via PCR any by-product transcript generated from exon skipping events (data not shown). Nevertheless, a few studies have observed a lack of correlation between expression patterns of parental and circRNA [11, 12]. The disproportionate expression patterns of circRNA may result from their accumulation over time due to their structural stability. However, active processing of circRNA from parental transcripts with the aid of RNA binding proteins such as DHX9, Quaking and Mbln are also contributing to their expression[12, 14, 29]. As some circRNAs are derived from novel exons of parental transcripts, we have demonstrated the importance of using circRNA software, such as Ularcirc, that does not rely on existing gene models.

Active circRNA biogenesis raises questions relating to their biological function. We incorporated into Ularcirc two basic functional analysis tools, ORF and miRNA motif analysis, that enable the screening for potential circRNA function. Ularcirc reconstructs circRNA sequences from genomic and transcriptional database coordinates, and the output of the functional analysis is represented as circos plots. We were surprised to identify circRNAs that have an ORF longer than the coding potential of an equivalent linear transcript. Hipk3 circRNA is particularly intruiging as it is present in the cytoplasm of many cell types and has been identified to act as a miRNA sponge [16]. Silencing of Hipk3 circRNA significantly inhibited cell growth which was attributed to miRNA sponge activity. However, with recent studies identifying that circRNAs can be translated reveal that Hipk3 circRNA function may also be explained by the translation of a novel isoform from its super-ORF. Also, under this scenario, the miRNA binding capabilities may be a translational regulatory event rather than pure sponge activity.

In this study we have highlighted the efficacy of integrating the analysis of FSJ and BSJs to analyse circRNAs. Ularcirc is specifically developed for this task for researchers with minimal bioinformatic skill to perform complex analysis by incorporating integrated visualisation combined with a friendly graphical user interface. Visualising complex data sets will enable faster processing and appreciation of data trends than scrutinizing data in tabulated formats and reports. Our integrative analysis of FSJ and BSJ using Ularcirc has already gleaned new insights into circRNA biology. Ularcirc is a valuable technical resource that will significantly enhance the analytical capacity for circRNA research.

## MATERIAL AND METHODS

### Classification of read reads aligned to circRNA

We define four circRNA alignments. Type I alignments are indistinguishable to alignments that are derived from linear RNA transcripts. Type II alignments capture BSJ in the first read pair, while type III alignments capture BSJ in the second read pair. Type IV alignments are discordant read pairs which flank a potential BSJ site.

### Distribution of BSJ reads

The proportion of type I alignments can be calculated using the formula:

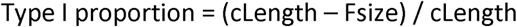

Where cLength is circRNA length and Fsize is library fragment size. The following R-code was used to generate figure 1C:

~~~
circRNA_Length <- 300:3000
Frag_Size <- 300
plot (circRNA_Length, y = (circRNA_Length - Frag_Size)/circRNA_Length * 100,
ylim=c(0,100))
~~~

The following R-code was used to generate figure 1D

~~~
circRNA_Length <- 300:3000
Frag_Size <- 300
Proportion_Type_II_III_IV <- (Frag_Size/circRNA_Length*100)
Proportion_Type_II_III <- Proportion_Type_II_III_IV/3*2
Proportion_Type_I <- 100- Proportion_Type_II_III_IV
BSJ_Reads <- 10
TypeI_Reads <- BSJ_Reads /Proportion_Type_II_III * 100
plot (circRNA_Length, TypeI_Reads)
~~~

### Implementation of Ularcirc

Ularcirc is written in R using the shiny framework. There are numerous functional aspects built into Ularcirc, below are descriptions of how these were implemented.

CircRNA sequence assembly: Sequences are only retrieved for backsplice junctions that overlap exactly with defined canonical exon boundaries of at least one transcript. In situations of genes that have multiple transcripts whose exons align to a BSJ, Ularcirc selects the transcript that will produce the longest sequence. Exon sequences are retrieved from the genome databases and are concatenated together.

Backsplice sequence assembly: The same process as for total circRNA sequence assembly however 50nt of sequence each side of the backspice junction is displayed. The backsplice junction is highlighted by a ‘.’ Character.

ORF detection: Open reading frames were calculated from circRNA sequence assembly and concatenating the total sequence for a total of three times. The longest open reading frames as calculated by the “translate” function from biostrings is displayed.

miRNA detection:Ularcirc queries miRbase.db bioconductor database and extracts the reverse complement of miRNA seed sequences. Seed sequences are matched against the selected circRNA candidate.

RAD score: Type II and type III alignments are determined from the CIGAR string of each sequence read that aligns to a specific backsplice junction within the chimeric read junction file. The RAD score is the ratio of type II alignments as calculated by the following formula:

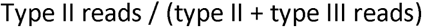

The RAD score is only calculated on backsplice junctions that have a raw count greater than 10.

CircRNA annotation: This is an optional feature. When selected all BSJ are annotated with overlapping potential parental genes names. Importantly there is no requirement for BSJ to overlap with exon boundaries.

### Parental and putative by-product transcript analysis

We define a circRNA parental transcript as a linear transcript whose exonic sequences overlap that of a identified circRNA. We do not exclude circRNAs that do not align perfectly to existing annotated exonic boundaries.

We define a parental by-product transcript as a linear transcript that contains a splice junction that excludes all sequences that give rise to a circRNA. This excluding splice junction does not have to be the exact coordinates of the backsplice junction.

Ularcirc is capable of building annotated tables that assembles raw counts of BSJ and FSJ. When selected the FSJ can be assigned to those that reside either (1) within the boundaries of a BSJ (i.e. representing forward splicing within circRNA or parental transcript), (2) external to that of a BSJ (forward splicing only to that of the parental transcript or (3) spanning a BSJ (forward splicing of a by-product transcript).

### Comparing Ularcirc to circExplorer2

The outputs of both Ularcirc and circExporer2 were compared using custom scripts. Each circRNA was identified by genomic coordinates of the backsplice junctions. For some circRNAs, circExplorer shifts BSJ coordinates in a realignment step to match exon junctions of the provided gene model. To ensure these candidates are correctly matched to Ularcirc entries, the genomic coordinates of the uniquely identified BSJ was compared across a 10nt window. If a corresponding BSJ from Ularcirc data was found, it was considered a match. The R code for this match process is provided in the Ularcirc package via the function Plot_Ularcirc_vs_circExplorer.

### Public Datasets

All public available datasets analysed in this study (listed in **supplementary table 2)** were downloaded from NCBI sequence read archives using sratoolkit.2.8.2 fastq-dump.

### Alignment

All datasets were trimmed for adaptor sequence and poor quality sequence using trimmomatic using the following options LEADING:3 TRAILING:3 SLIDINGWINDOW:4:15 MINLEN:33. All data sets were aligned with the STAR aligner against either hg38 or mm10 genome using the two Pass star alignment protocol [34]. On the second pass alignment the following STAR options were set to capture chimeric reads. After alignment the chimeric output file and SJ.out files were renamed to have a common unique prefix for each sample and imported into Ularcirc for downstream analysis.

### Generated datasets

For the dataset of murine epithelial cells of the small intestine, we used 1 μg of total RNA extracted from wild type adult mouse as in [35] and generated the NGS library prep using the “TruSeq stranded total RNA sample preparation” kit from Illumina strictly following manufacturer instructions (with depletion of ribosomal RNA using the “RiboZero rRNA removal mix”). Library was sequenced using an Illumina 2500 and fastq files were provided by the precinct sequencing provider.

### RT-PCR

Total RNA was obtained from epithelial cells of the small intestine isolated as described in [35] or from small intestine of human biopsies (Clontech). 1 ug of total RNA was reverse transcribed using the SuperScript III First-Strand Kit (Invitrogen) and following manufacturer instructions. 1 or 2 uL of cDNA (undiluted or diluted 1:10) was used as input for PCR amplification with Biomix Red (Bioline). Conditions of PCR are 95°C for 30 s, 60°C for 30 s and 72°C for 30 s for 30 or 35 cycles. The PCR products were visualized on TAE 2% agarose gel.

Sequence of *ApoA4* primers used in Figure 6 are listed below:

- a mouse: 5’-CCTGCAGTCAATCTGCACA
- b mouse: 5-AGTGACATCCGTCTTCTGAA
- c mouse: 5’-GATGCTAGTACGTATGCTGA
- b mouse for HTS: 5-CCTCTCTATGGGCAGTCGGTGATAGTGACATCCGTCTTCTGAA
- c mouse for HTS: 5’-CCATCTCATCCCTGCGTGTCTCCGACTCAGGATGCTAGTACGTATGCTGA
- d mouse: 5’-GCTTAGCTGGGTAAACGTAG (15 first nt complementary to region in exon 2; last 5 nt complementary to region in exon 3)
- a human: 5-TGTAGGGAGGATCCAGTGT
- b human (for PCR with a human): 5’-CCTGGTCAGCACTGACCT
- b human (for PCR with c human): 5’-AGTTGCTGGGTGAGTTCAGA
- c human: 5’-TCTTCCAGGACAAACTTGGA

### Sanger Sequencing

Products i and ii amplified by PCR Figure 6E were isolated and purified from gel with Wizard kit (Promega) and send for Sanger sequencing with primer c mouse: 5’-GATGCTAGTACGTATGCTGA to the Australian Genome Research Facility (AGRF).

## DECLARATIONS

## Acknowledgements

All animal experimentation was approved by the Animal Ethics Committee of the Children’s Medical Research Institute and of the Children’s Hospital at Westmead, Sydney, Australia. We thank the CMRI BioServices Unit for animal care.

## Availability of data and materials

Ularcirc software is available at https://github.com/VCCRI/Ularcirc, which is under GNU general public licencse v3.0. The public RNA-Seq datasets used in this study are listed in supplementary table 1. The generated RNA-seq data sets can be accessed via the SRA

Operating system: Platform independent

Programming language: R

Other requirements: See Github page

## Authors’ contributions

DTH conceived the study, designed the pipeline, implemented the program, and wrote the manuscript. NF generated the murine small intestine datasets with the help of DTH, conducted experimental validation of ApoA4 circRNAs and helped with manuscript and designing figures. PPLT contributed resources for the murine small intestine datasets and for ApoA4 experiments, and contributed to the final manuscript. JWKH contributed to project design and final manuscript. All authors read and approved the final manuscript.

## Competing interests

The authors declare that they have no competing interests.

Ethics approval and consent to participate

Not applicable.

## References

1. Hossain ST, Malhotra A, Deutscher MP: How RNase R Degrades Structured RNA: ROLE OF THE HELICASE ACTIVITY AND THE S1 DOMAIN. J Biol Chem 2016, 291: 7877–7887.

2. Jeck WR, Sharpless NE: Detecting and characterizing circular RNAs. Nat Biotechnol 2014, 32: 453–461.

3. Hansen TB, Jensen TI, Clausen BH, Bramsen JB, Finsen B, Damgaard CK, Kjems J: Natural RNA circles function as efficient microRNA sponges. Nature 2013, 495: 384–388.

4. Jeck WR, Sorrentino JA, Wang K, Slevin MK, Burd CE, Liu J, Marzluff WF, Sharpless NE: Circular RNAs are abundant, conserved, and associated with ALU repeats. RNA 2013,19:141–157.

5. Memczak S, Jens M, Elefsinioti A, Torti F, Krueger J, Rybak A, Maier L, Mackowiak SD, Gregersen LH, Munschauer M, et al: Circular RNAs are a large class of animal RNAs with regulatory potency. Nature 2013, 495: 333–338.

6. Chu Q, Zhang X, Zhu X, Liu C, Mao L, Ye C, Zhu QH, Fan L: PlantcircBase: A Database for Plant Circular RNAs. Mol Plant 2017, 10: 1126–1128.

7. Glazar P, Papavasileiou P, Rajewsky N: circBase: a database for circular RNAs. RNA 2014, 20: 1666–1670.

8. Ebbesen KK, Kjems J, Hansen TB: Circular RNAs: Identification, biogenesis and function. Biochim Biophys Acta 2016, 1859: 163–168.

9. Salzman J, Gawad C, Wang PL, Lacayo N, Brown PO: Circular RNAs are the predominant transcript isoform from hundreds of human genes in diverse cell types. PLoS One 2012, 7:e30733.

10. Zhang XO, Wang HB, Zhang Y, Lu X, Chen LL, Yang L: Complementary sequence-mediated exon circularization. Cell 2014,159:134–147.

11. Rybak-Wolf A, Stottmeister C, Glazar P, Jens M, Pino N, Giusti S, Hanan M, Behm M, Bartok O, Ashwal-Fluss R, et al: Circular RNAs in the Mammalian Brain Are Highly Abundant, Conserved, and Dynamically Expressed. Mol Cell 2015, 58: 870–885.

12. Conn SJ, Pillman KA, Toubia J, Conn VM, Salmanidis M, Phillips CA, Roslan S, Schreiber AW, Gregory PA, Goodall GJ: The RNA binding protein quaking regulates formation of circRNAs. Cell 2015, 160: 1125–1134.

13. Khan MA, Reckman YJ, Aufiero S, van den Hoogenhof MM, van der Made I, Beqqali A, Koolbergen DR, Rasmussen TB, van der Velden J, Creemers EE, Pinto YM: RBM20 Regulates Circular RNA Production From the Titin Gene. Circ Res 2016,119:996–1003.

14. Ashwal-Fluss R, Meyer M, Pamudurti NR, Ivanov A, Bartok O, Hanan M, Evantal N, Memczak S, Rajewsky N, Kadener S: circRNA biogenesis competes with pre-mRNA splicing. Mol Cell 2014, 56: 55–66.

15. Ivanov A, Memczak S, Wyler E, Torti F, Porath HT, Orejuela MR, Piechotta M, Levanon EY, Landthaler M, Dieterich C, Rajewsky N: Analysis of intron sequences reveals hallmarks of circular RNA biogenesis in animals. Cell Rep 2015,10:170–177.

16. Zheng Q, Bao C, Guo W, Li S, Chen J, Chen B, Luo Y, Lyu D, Li Y, Shi G, et al: Circular RNA profiling reveals an abundant circHIPK3 that regulates cell growth by sponging multiple miRNAs. Nat Commun 2016, 7: 11215.

17. Holdt LM, Stahringer A, Sass K, Pichler G, Kulak NA, Wilfert W, Kohlmaier A, Herbst A, Northoff BH, Nicolaou A, et al: Circular non-coding RNA ANRIL modulates ribosomal RNA maturation and atherosclerosis in humans. Nat Commun 2016, 7: 12429.

18. Pamudurti NR, Bartok O, Jens M, Ashwal-Fluss R, Stottmeister C, Ruhe L, Hanan M, Wyler E, Perez-Hernandez D, Ramberger E, et al: Translation of CircRNAs. Mol Cell 2017, 66:9–21 e27.

19. Yang Y, Fan X, Mao M, Song X, Wu P, Zhang Y, Jin Y, Yang Y, Chen LL, Wang Y, et al: Extensive translation of circular RNAs driven by N6-methyladenosine. Cell Res 2017, 27: 626–641.

20. Hansen TB, Veno MT, Damgaard CK, Kjems J: Comparison of circular RNA prediction tools. Nucleic Acids Res 2016, 44:e58.

21. Gao Y, Wang J, Zheng Y, Zhang J, Chen S, Zhao F: Comprehensive identification of internal structure and alternative splicing events in circular RNAs. Nat Commun 2016, 7: 12060.

22. Engstrom PG, Steijger T, Sipos B, Grant GR, Kahles A, Ratsch G, Goldman N, Hubbard TJ, Harrow J, Guigo R, et al: Systematic evaluation of spliced alignment programs for RNA-seq data. Nat Methods 2013, 10: 1185–1191.

23. Baruzzo G, Hayer KE, Kim EJ, Di Camillo B, FitzGerald GA, Grant GR: Simulation-based comprehensive benchmarking of RNA-seq aligners. Nat Methods 2017,14:135–139.

24. Phanstiel DH, Boyle AP, Araya CL, Snyder MP: Sushi.R: flexible, quantitative and integrative genomic visualizations for publication-quality multi-panel figures. Bioinformatics 2014, 30: 2808–2810.

25. Szabo L, Morey R, Palpant NJ, Wang PL, Afari N, Jiang C, Parast MM, Murry CE, Laurent LC, Salzman J: Statistically based splicing detection reveals neural enrichment and tissue-specific induction of circular RNA during human fetal development. Genome Biol 2015,16:126.

26. Li XF, Lytton J: A circularized sodium-calcium exchanger exon 2 transcript. J Biol Chem 1999, 274: 8153–8160.

27. Giudice J, Xia Z, Wang ET, Scavuzzo MA, Ward AJ, Kalsotra A, Wang W, Wehrens XH, Burge CB, Li W, Cooper TA: Alternative splicing regulates vesicular trafficking genes in cardiomyocytes during postnatal heart development. Nat Commun 2014, 5: 3603.

28. Aufiero S, van den Hoogenhof MMG, Reckman YJ, Beqqali A, van der Made I, Kluin J, Khan MAF, Pinto YM, Creemers EE: Cardiac circRNAs arise mainly from constitutive exons rather than alternatively spliced exons. RNA 2018.

29. Aktas T, Avsar Ilik I, Maticzka D, Bhardwaj V, Pessoa Rodrigues C, Mittler G, Manke T, Backofen R, Akhtar A: DHX9 suppresses RNA processing defects originating from the Alu invasion of the human genome. Nature 2017, 544: 115–119.

30. Wong WM, Gerry AB, Putt W, Roberts JL, Weinberg RB, Humphries SE, Leake DS, Talmud PJ: Common variants of apolipoprotein A-IV differ in their ability to inhibit low density lipoprotein oxidation. Atherosclerosis 2007,192:266–274.

31. Barrett SP, Wang PL, Salzman J: Circular RNA biogenesis can proceed through an exon-containing lariat precursor. Elife 2015, 4:e07540.

32. Guarnerio J, Bezzi M, Jeong JC, Paffenholz SV, Berry K, Naldini MM, Lo-Coco F, Tay Y, Beck AH, Pandolfi PP: Oncogenic Role of Fusion-circRNAs Derived from Cancer-Associated Chromosomal Translocations. Cell 2016,165:289–302.

33. Liang D, Tatomer DC, Luo Z, Wu H, Yang L, Chen LL, Cherry S, Wilusz JE: The Output of Protein-Coding Genes Shifts to Circular RNAs When the Pre-mRNA Processing Machinery Is Limiting. Mol Cell 2017, 68:940–954 e943.

34. Dobin A, Davis CA, Schlesinger F, Drenkow J, Zaleski C, Jha S, Batut P, Chaisson M, Gingeras TR: STAR: ultrafast universal RNA-seq aligner. Bioinformatics 2013, 29: 15–21.

35. Fossat N, Tourle K, Radziewic T, Barratt K, Liebhold D, Studdert JB, Power M, Jones V, Loebel DA, Tam PP: C to U RNA editing mediated by APOBEC1 requires RNA-binding protein RBM47. EMBO Rep 2014, 15: 903–910.

